# A tradeoff between tolerance and resistance to a major fungal pathogen in elite wheat cultivars

**DOI:** 10.1101/354357

**Authors:** Alexey Mikaberidze, Bruce A. McDonald

**Affiliations:** Plant Pathology, Institute of Integrative Biology, ETH Zurich

**Keywords:** *Triticum aestivum*, host-pathogen interaction, host defenses, plant disease, *Zymoseptoria tritici*, digital phenotyping

## Abstract

- Tolerance and resistance represent two strategies that hosts evolved to protect themselves from pathogens. Tolerance alleviates the reduction in host fitness due to infection without reducing a pathogen’s growth, while resistance reduces pathogen growth. We investigated tolerance of wheat to the major fungal pathogen *Zymoseptoria tritici* in 335 elite wheat cultivars.
- We used a novel digital phenotyping approach that included 11,152 infected leaves and counted 2,069,048 pathogen fruiting bodies.
- We discovered a new component of tolerance that is based on the relationship between the green area remaining on a leaf and the number of pathogen fruiting bodies. We found a negative correlation between tolerance and resistance among intolerant cultivars, presenting the first compelling evidence for a tradeoff between tolerance and resistance to plant pathogens. Surprisingly, the tradeoff arises due to limits in the host resources available to the pathogen and not due to metabolic constraints, contrary to what ecological theory suggests.
- The mechanism underlying this tradeoff may be relevant for many plant diseases in which the amount of host resources available to the pathogen can limit the pathogen population. Our analysis indicates that European wheat breeders may have selected for tolerance instead of resistance to an important pathogen.

## Introduction

Tolerance and resistance represent two important mechanisms that plants and animals evolved to protect themselves from pathogens (Roy et al., 2000; Baucom & De Roode, 2011). Tolerance to a pathogen is usually defined as the host’s ability to alleviate the reduction in its fitness due to infection without reducing the growth of the pathogen (Baucom & De Roode, 2011; Ney et al., 2013). A more tolerant host genotype will suffer a smaller loss in fitness per unit increase of pathogen population present within the host (called the pathogen burden) than a less tolerant host genotype. Hence, tolerance of a host genotype can be quantified as the reduction in host fitness per unit increase in pathogen burden. In contrast, resistance is usually measured as the host’s ability to suppress the infection itself and reduce the resulting pathogen burden upon infection. The difference between tolerance and resistance is illustrated in Fig. 1.

**Figure 1.**
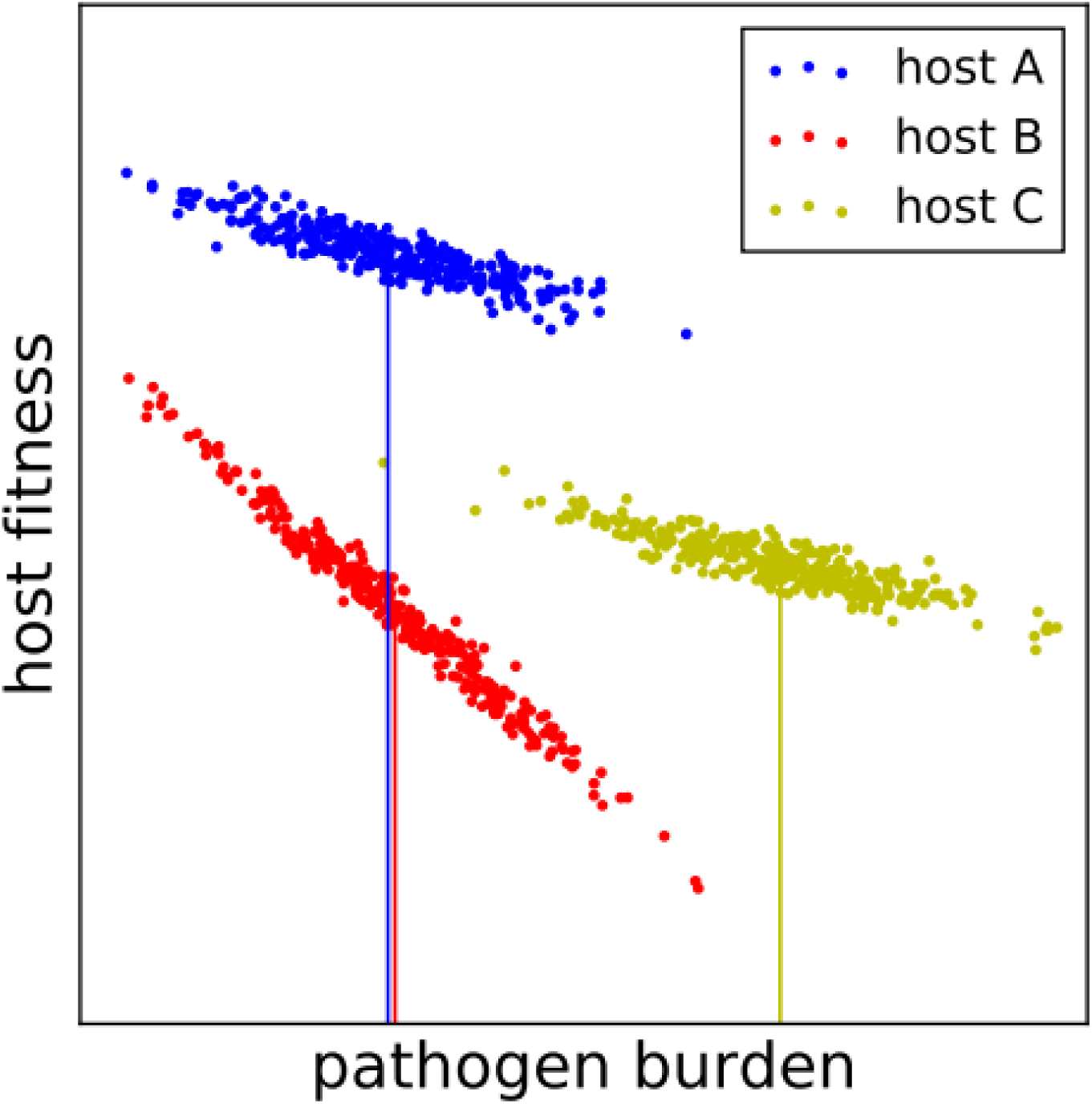
Differentiation and quantification of tolerance and resistance. The figure illustrates the relationships between host fitness and pathogen load for three hypothetical host genotypes: A (blue), B (red) and C (yellow). For each host genotype, tolerance is quantified as the rate at which the host loses its fitness with an increase in pathogen burden. The difference in resistance between hosts can be measured on the X-axis as the difference in the mean pathogen burden (vertical lines). The position of each genotype on the Y-axis is determined by its fitness in the absence of the pathogen and is not related to tolerance or resistance. Host genotypes A and B have the same resistance because the average pathogen burden that they carry is the same. However, genotype A is more tolerant than genotype B because the fitness of genotype B decreases at a higher rate with increasing pathogen burden. This is reflected in the steeper slope in genotype B compared to genotype A. In contrast, host genotypes A and C have the same tolerance, but genotype A is more resistant than genotype C. Finally, the comparison of host genotype B and host genotype C represents a mixed case: genotype B is more resistant, but less tolerant than genotype C.

Since the concept of tolerance was first coined more than a century ago (Cobb, 1894), numerous studies [reviewed by Pagan & Garcia-Arenal (2018)] investigated tolerance to pathogens in crop plants (Caldwell *et al.* 1958; Schafer, 1971; Newton *et al.*, 1998; Bingham *et al.*, 2009; Ney *et al.*, 2013; Newton, 2016), model plants (Kover & Schaal, 2002; Pagan et al., 2008, Shuckla et al., 2017) and wild plants (Roy et al., 2000; Inglese & Paul, 2006; Carr et al., 2006). Råberg et al. (2007) were the first to demonstrate tolerance to an infectious disease in animals. The therapeutic potential of tolerance in human medicine inspired a surge of further investigations (Medzhitov et al., 2012; Ayres & Schneider, 2012; Råberg, 2014; Soares et al., 2017). Several studies uncovered molecular mechanisms of tolerance in model animal species (Ayres & Schneider, 2008; Shinzawa et al., 2009; Richardson et al., 2010; Maze-Guilmo et al., 2014), while Blanchet et al. (2010); Jackson et al. (2014); Hayward et al. (2014); Maze-Guilmo et al. (2014) characterized tolerance to parasites in wild animal populations. Zeller & Koella (2017) used an experimental evolution approach to determine how tolerance/resistance evolves in mosquito populations exposed to microsporidian parasites.

It is generally thought that since both tolerance and resistance are defense strategies that require reallocation of host resources, they should confer fitness costs to the host (Roy & Kirchner, 2000; Simms & Triplett, 1994; Brown, 2002). For this reason, a metabolic tradeoff between tolerance and resistance is expected due to a limitation in host resources. A large body of ecological theory has been developed based on this premise (van der Meijden et al., 1988; Herms & Mattson, 1992; Roy & Kirchner, 2000; Fornoni et al., 2004; Restif & Koella, 2004; Miller et al., 2005; Best et al., 2008). However, empirical evidence for a tradeoff between tolerance and resistance remains sparse. A few studies reported a negative relationship between tolerance and resistance to herbivory (Fineblum & Rausher, 1995; Stowe, 1998; Baucom & Mauricio, 2008) and Råberg et al. (2007) presented a similar finding in mice infected with malaria. Other studies reported no correlation between tolerance and resistance in plants subjected to herbivores (Mauricio et al., 1997), in humans infected with HIV (Regoes et al., 2014) or in wild sheep infected with a parasite (Maze-Guilmo et al., 2014). Interestingly, in *Drosophila melanogaster* populations exposed to a bacterial infection, tolerance and resistance exhibited a positive correlation (Howick & Lazzaro, 2014). Likewise, populations of the mosquito *Aedes aegypti* that were infected by the microsporidian parasite *Vavraia culicis* and evolved for 10 generations exhibited a positive relationship between tolerance and resistance (Zeller & Koella, 2017). No evidence for a tradeoff between host tolerance and resistance was so far reported in the plant pathology literature.

In this study, we investigated tolerance to the fungal pathogen *Zymoseptoria tritici* (formerly *Mycosphaerella graminicola*) in 335 elite European wheat cultivars. *Z. tritici* causes septoria tritici blotch (STB), a disease that is a major constraint on wheat production globally and the most destructive disease of wheat in Europe (Fones & Gurr, 2015). *Z. tritici* spores germinate on wheat leaves and penetrate the leaves through stomata (Kema et al., 1996). After penetration, the fungus grows for several days within leaves without producing visible symptoms. During this asymptomatic period, the pathogen invades the host mesophyll around the position of the initial penetration. After 10 to 20 days of asymptomatic growth, the fungus becomes necrotrophic and kills the invaded plant tissue, forming necrotic lesions. Asexual fruiting bodies called pycnidia begin to form in the necrotic lesions soon thereafter. Spores that form in the pycnidia provide inoculum to start the next cycle of pathogen reproduction. The formation of necrotic lesions corresponds to host damage caused by the pathogen that can be quantified as the proportion of leaf area covered by lesions (PLACL). The number of pycnidia provides a measure of pathogen reproduction that can be quantified by counting the number of pycnidia present on an infected leaf, *N*_*p*_ (Stewart et al., 2016a; Karisto et al., 2018).

Control of STB relies mainly on applications of fungicides and deployment of STB-resistant wheat varieties. However, populations of *Z. tritici* are extremely diverse due to a high degree of sexual reproduction and large effective population sizes. As a result, the pathogen has the capacity to rapidly adapt to both fungicides (Fraaije et al., 2005; Zhan et al., 2006) and host resistances (Cowger et al., 2000; McDonald and Mundt, 2016) as a result of strong directional selection favoring particular pathogen genotypes. In contrast, host tolerance does not impair pathogen reproduction and is not expected to impose strong directional selection. For this reason, tolerance presents a promising alternative to protect wheat yield that is not prone to pathogen adaptation.

Several previous studies investigated tolerance of wheat to STB empirically (Eyal & Ziv, 1974; Zuckerman et al., 1997; Parker et al., 2004; Foulkes et al., 2006; Collin et al., 2018). Van den Berg et al (2017) used mathematical modeling to reveal functional traits in wheat that contribute to tolerance. These studies used wheat yield (measured as tons of grain per hectare or as the thousand kernel weight) to quantify the plant fitness (the Y-axis in Fig. 1) and the PLACL or healthy area duration (HAD, Waggoner & Berger (1987)) to quantify the pathogen burden (the X-axis in Fig. 1). Accordingly, tolerance was quantified as the yield loss associated with each unit increase in PLACL or unit loss in HAD.

PLACL and HAD quantify the damage that the pathogen causes on an infected host plant. However, these quantities do not necessarily accurately reflect the size of the pathogen population present within the infected host plant (Stewart et al., 2016a; Karisto et al., 2018). For this reason, tolerance measured in these traditional ways is considered to be tolerance to the disease, which may not coincide with tolerance to the pathogen (Gaunt, 1981). The goal in this study was to characterize wheat tolerance to its pathogen, *Z. tritici*. With this in mind, we used (i) green leaf area to quantify a component of plant fitness and (ii) the number of pycnidia per leaf to quantify the pathogen burden. Grain yield is usually seen as a more comprehensive measure of fitness in crop plants than green leaf area. However, a number of field experiments have demonstrated that the reduction in the green area of the three upper-most leaf layers in wheat is a major driver of yield loss induced by STB (Eyal & Ziv, 1974; King et al., 1983; Forrer & Zadoks, 1983; Shaw & Royle 1989b, Thomas et al., 1989), thereby justifying our choice (i) (see Discussion for a more detailed justification). The choice (ii) is justified because the number of pycnidia per leaf was shown to be the main factor influencing the number of pathogen spores produced on an infected leaf (Stewart et al., 2016a). Moreover, the proportion of the leaf area covered by STB lesions was demonstrated to be largely independent from the number of pycnidia produced on a leaf (Karisto et al., 2018). For these reasons, the number of pycnidia per leaf is a better indicator of the pathogen population inhabiting a leaf than the PLACL.

By conducting these measurements on 11,152 individual wheat leaves belonging to 335 different cultivars (counting in total 2,069,048 individual pycnidia), we were able to identify and measure a novel component of wheat tolerance to *Z. tritici* that operates on the scale of individual leaves. We call this “leaf tolerance” as opposed to the “whole-plant tolerance” that was characterized previously. In this study, we focused on leaf tolerance and did not consider whole-plant tolerance. A way to estimate tolerance over a range of pathogen burdens as we describe here is to estimate range tolerance (Baucom & De Roode, 2011). The component of tolerance that we measured represents fecundity tolerance rather than mortality tolerance, because this disease does not kill its host but instead reduces its fecundity.

We used a combination of mathematical modeling and field experimentation to formulate and test several hypotheses connected to leaf tolerance of wheat to *Z. tritici*. First, based on our current understanding of the infection biology of *Z. tritici*, we formulated and tested empirically two alternative hypotheses regarding the relationship between the green leaf area and the number of pycnidia per leaf. Second, we tested the hypothesis that wheat cultivars differ in terms of their leaf tolerance. Finally, we tested the expectation of a tradeoff between leaf tolerance and resistance and found a significant negative relationship between tolerance and resistance. Surprisingly, our analysis indicates that this negative association arises due to the limitation in the leaf area of wheat plants and not as a result of metabolic costs associated with tolerance/resistance as predicted by ecological theory.

## Materials and Methods

Here we analyzed a subset of the raw data reported in (Karisto et al., 2018). Below, we describe the main features of the experimental design that are relevant for this analysis. A comprehensive description of the experimental design can be found in (Karisto et al., 2018).

### Plant materials and experimental design

In total, 335 elite European winter wheat (*Triticum aestivum*) varieties from the GABI-wheat panel (Kollers et al., 2013a,b) were evaluated in this experiment. Two biological replicates of the wheat panel were grown during the 2015-16 growing season in two complete blocks separated by approximately 100 m at the Field Phenotyping Platform site of the Eschikon Field Station of the ETH Zurich, Switzerland (coordinates 47.449°N, 8.682°E) (Kirchgessner et al., 2017). The complete blocks were composed of 18 rows and 20 columns consisting of 1.2 x 1.7 m plots, with the genotypes arranged randomly within each block. Best practices recommended for conventional, high-input wheat production were used, including applications of fertilizers and pesticides. Complete details are given in (Karisto et al., 2018).

### Septoria tritici blotch inoculum, sampling of infected leaves

All STB infection was natural, with the majority of primary inoculum likely originating from airborne ascospores coming from nearby wheat fields that surround the Eschikon field site. As a result, the infections analyzed in this experiment were caused by thousands of different pathogen strains. For this study we used leaves exhibiting obvious STB lesions that were collected on 4 July 2016 (approximate range of GS 75 [milk development] to GS 85 [dough development]). Up to 16 infected leaves were collected at random for each plot from the leaf layer below the flag leaf (i.e., flag-1 or second leaf). The sampled leaves were placed into paper envelopes, kept on ice in the field, and stored at 4°C for 2 days before mounting onto A4 paper with printed reference marks and sample names, as described by Stewart et al. (2016b). Absorbent paper was placed between each sheet of eight mounted leaves and sheets were pressed with approximately 5 kg at 4°C for 2 to 3 days prior to scanning at 1,200 dpi with a Canon CanoScan LiDE 220 flatbed scanner.

### Determination of the green leaf area and the number of pycnidia per leaf

Scanned images were analyzed with the software ImageJ (Schindelin et al., 2015) using the macro described by Karisto et al. (2018). The maximum length of the scanned area for each leaf was 17 cm. When leaves were longer than 17 cm, bases of the leaves were placed within the scanned area, while the leaf tips extended outside the scanned area. For each leaf, the following quantities were automatically recorded from the scanned image: total leaf area (*A*_tot_), necrotic and chlorotic leaf area (*A*_necr_) and the number of pycnidia (*N*_p_). Necrotic and chlorotic leaf areas were detected based on discoloration of the leaf surface and were not based on the presence of pycnidia. We then calculated the green (healthy) leaf area as *H* = *A*_tot_ -*A*_necr_.

### Statistical analysis

Statistical analysis was conducted in the Python programming language (version 3.6.2, https://www.python.org) using the open-source packages scipy (version 0.19.1), numpy (version 1.11.1) and matplotlib (version 1.5.3; Jones et al., 2001). The Python package rpy2 (version 2.8.6, https://rpy2.bitbucket.io/) was used to access statistical routines of R (R Core Team, 2016).

To control for the effect of total leaf area on the number of pycnidia per leaf, we performed the adjustment *N*_p,i_ →(*A*_tot_ /*A*_tot,i_) *N*_p,i_ prior to the analysis, where *N*_p,i_ and *A*_tot,i_ is the number of pycnidia and the total area of an individual leaf *i* and *A*_tot_ is the mean total leaf area averaged over the whole dataset. First, we pooled together the data from leaves belonging to different cultivars and fitted the relationship between *N*_*p*_ and *H* using a linear function *H* =*H*_0_ (1 −*κ N*_*p*_) and an exponential function *H* =*H* _0_ exp(− *κ N*_*p*_), where *H* is the green leaf area and *H*_*0*_ is the green leaf area in the absence of disease. For both functions, the slope *κ* can be used to quantify tolerance: small *κ*-values correspond to high leaf tolerance and large *κ*-values correspond to low leaf tolerance. This overall fit gave us the baseline to which we then compared the tolerance of individual wheat cultivars. Second, we estimated *κ* in each of the 335 wheat cultivars by fitting both the linear and the exponential functions to individual leaf data belonging to each of the cultivars. Next, we used multiple one-sided bootstrap t-tests with resampling cases (Davison and Hinkley, 2001) to compare *κ*-estimates in each cultivar to the baseline *κ*-estimate, where we used the false discovery rate correction for multiple comparisons. Fits were performed using the nonlinear ordinary least-squares optimization with the Nelder-Mead method in the lmfit package (version 0.9.7) for Python (Newville et al., 2014). We also determined the significance of the effects of the spatial block and the cultivar on tolerance: we used likelihood ratio tests to compare more complex models in which data from each cultivar/spatial block was fitted using separate *κ* or *H* _0_-values to simpler models where only single *κ* or *H* _0_ parameters were fitted to the whole dataset.

To determine whether there is a relationship between tolerance and resistance of wheat to *Z. tritici*, we used the correlation test based on the Spearman’s rank correlation coefficient, *r*_S_ (routine “scipy.stats.spearmanr” of the scipy package for Python), to analyze correlations between tolerance quantified as the slope *κ* and resistance quantified as the average number of pycnidia per leaf (Karisto et al., 2018). We chose to use the Spearman’s correlation instead of Pearson’s correlation, because it is computed in a non-parametric fashion based on relative ranks of the estimates. It does not assume any specific functional form of the relationship between the two variables and thereby is not influenced by the widths of their distributions. To test whether the correlation is significantly different from zero, the routine uses a t-test that requires a number of assumptions to be fulfilled such as the normality of the probability distribution of the correlation coefficients.

To determine whether these assumptions hold and the conclusions based on this test are valid, we conducted a series of statistical tests based on a more robust bootstrap t-test (Davison et al., 1997). We generated a large number of bootstrap samples (*n*_bs_=10^5^) by resampling with replacement the estimates of leaf tolerance and resistance for each of the 335 cultivars and used them to compute the 95 % confidence interval (CI) of *r*_S_. We then generated the same number of bootstrap samples based on the estimates of tolerance and resistance separately in each of the two groups of cultivars, tolerant and intolerant cultivars. This allowed us to compute the confidence intervals of *r*_S_ estimates in each of the two cultivar groups. Finally, we tested whether *r*_S_ was significantly different from zero in each of the groups and whether one of the groups had a significantly higher *r*_S_ than the other group.

## Results

### A novel component of tolerance

Recent studies (Karisto et al., 2018; Stewart et al., 2016a) demonstrated that factors responsible for leaf damage during the infection are largely uncoupled from the pathogen’s capacity to reproduce on leaves. Based on this knowledge, we devised a simple mathematical model that describes the change in the green leaf area corresponding to an increase in the number of pycnidia on a leaf (for more details see Notes S1).

The following differential equation governs the relationship between the number of pycnidia and the green leaf area remaining on the leaf:

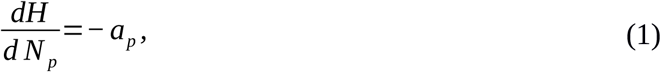

where *a*_*p*_ is the area of the lesion that corresponds on average to a single pycnidium. Equation (1) represents mathematically a rather general statement that the green area remaining on leaves decreases with increasing numbers of pycnidia. If *a*_*p*_ depends neither on *H*, nor on *N*_*p*_, then the solution of Eq. (1) is a linear function:

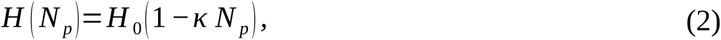

where *κ* =*a*_*p*_/*H*_0_ and *H*_0_ is the green leaf area in the absence of infection. Alternatively, if *a*_*p*_ is proportional to the green leaf area *H*, i.e. *a*_*p*_=*κH*, then the solution of Eq. (1) is an exponential function:

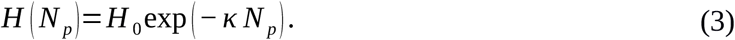

In both its linear and exponential versions, the model predicts that the leaf loses its green area as it carries higher numbers of pycnidia. We consider the green leaf area, *H*, as a quantity representing plant fitness and the number of pycnidia per leaf, *N*_*p*_, as a proxy for the pathogen population present on the leaf (pathogen burden). Consequently, the slope of the decrease, *κ*, characterizes the tolerance of wheat to *Z. tritici*: it measures the amount by which the green leaf area decreases when a single pycnidium is added to the leaf. We call *κ* the intolerance parameter, as the cultivars with higher *κ*-values will lose their green leaf area at a higher rate than cultivars with lower *κ*-values when the number of pycnidia is increased.

Using this model we developed a novel way to measure tolerance of wheat to *Z. tritici* that operates on the scale of individual leaves (“leaf tolerance”) as opposed to the “whole-plant tolerance” that was studied previously in this pathosystem (Eyal & Ziv, 1974; Parker et al., 2004; Foulkes et al., 2006; Collin et al., 2018). We demonstrated in Notes S2 that these two components of tolerance contribute to overall tolerance as multiplicative factors (under the assumption that the relationships between the yield and the damaged leaf area and between the yield and the number of pycnidia per leaf are both linear). Based on two sets of biological assumptions, we formulated two hypotheses about the shape of the relationship between the green leaf area and the number of pycnidia on the leaf represented by Eq. (2) and (3). We next tested these hypotheses using the empirical data gathered from wheat leaves naturally infected by *Z. tritici*.

### Relationship between green leaf area and yield

To determine whether the green leaf area in our experiment can be considered a good measure of plant fitness, we studied the correlation between the green leaf area measured on infected leaves and yield measured as the weight of grains per unit area of land [tons per hectare (t/ha)] and as the thousand kernel weight (TKW). Figure 2 illustrates the outcomes: green leaf area averaged over leaves belonging to the same cultivar correlates weakly, but significantly, with the yield of the corresponding cultivar (*r*_*S*_ =0.13, *p*=0.017 for yield measured in t/ha and *r*_*S*_ =0.13, *p*=0.019 for yield measured as TKW).

**Figure 2.**
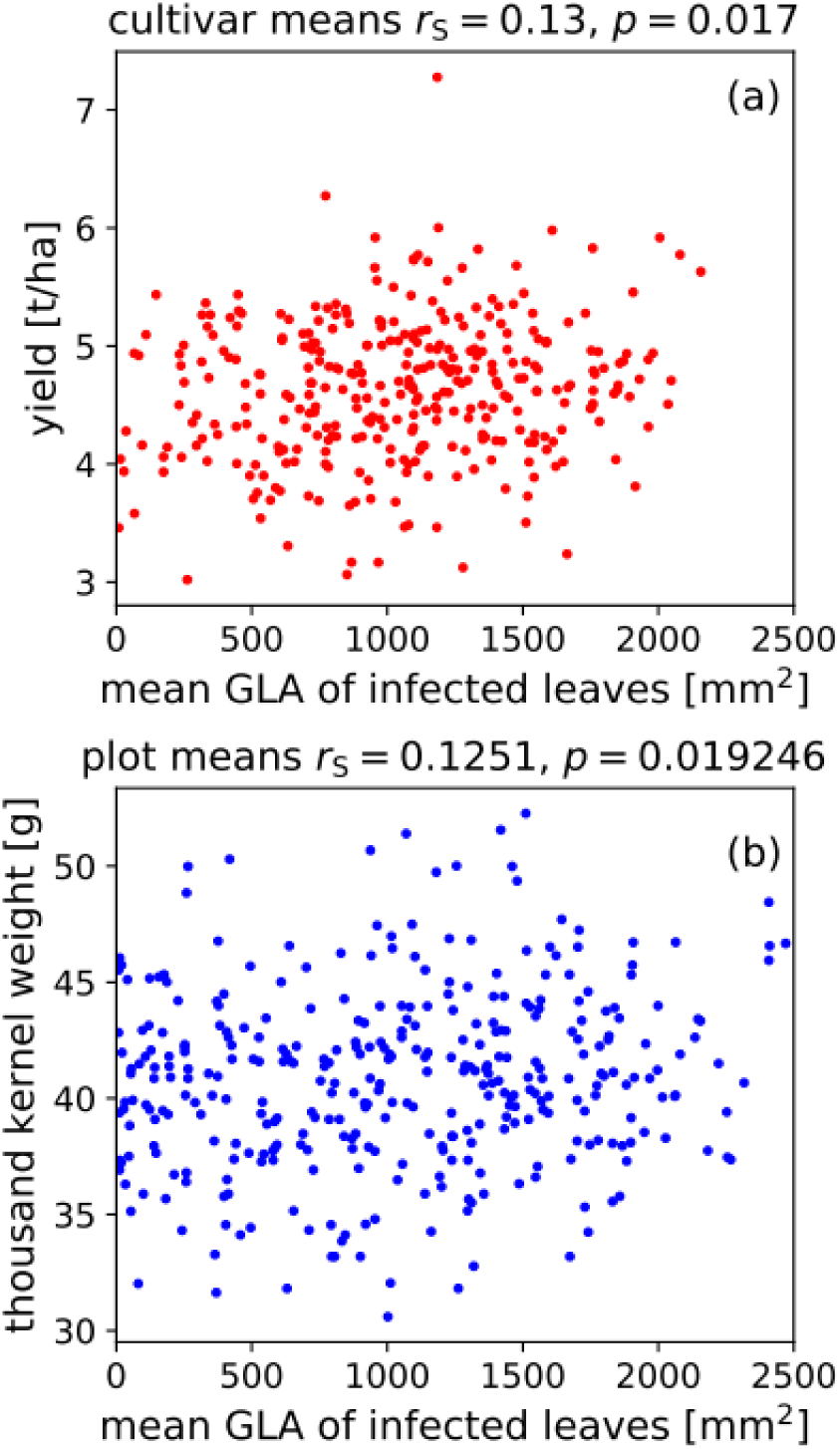
Correlation between the green leaf area (GLA) and wheat yield. (a) Yield in tons per hectare is plotted against the GLA of infected leaves measured at GS 75-85. Each value on the *x*-axis represents the average value over approximately 30 leaves originating from two different plots belonging to the same cultivar. Each value on the *y*-axis represents the yield averaged over two plots planted with the same cultivar. (b) Yield measured as thousand kernel weight is plotted against the GLA. Each value on the *x*-axis represents the average value over approximately 15 leaves originating from a single plot belonging to the same cultivar. Each value on the *y*-axis represents the yield measured in a single plot.

We would like to emphasize that in our experiment, the green leaf area was recorded only on infected leaves, while yield was measured from plants sampled without regard to their infection status, hence the yield measures comprised both healthy and infected plants. If in addition to the green area of infected leaves, we were able to also measure the STB incidence (that is the proportion of second leaves that were infected), then the product of the green leaf area on infected leaves times the STB incidence would give us the average green leaf area on all second leaves. This quantity would likely explain a much larger percentage of variation in yield. This has been convincingly demonstrated in a large number of field experiments, in which the reduction in wheat yield was strongly correlated with the reduction in the green leaf area of second leaves due to STB (e.g., King et al., 1983; Shaw & Royle, 1989b).

### Green leaf area decreases nonlinearly with the number of pycnidia

Figure 3a shows 11,152 individual leaf measurements of the number of pycnidia per leaf, *N*_*p*_, and the green leaf area, *H*. Overall, leaves lose more of their green area when they carry a larger number of pycnidia. The exponential function, Eq. (3), provided a better fit (standard error of the estimate *s*=472, coefficient of determination *R*^2^=0.37) than the linear function, Eq. (2) (*s*=508, *R*^2^=0.28). For this reason, we estimated the overall slope using the exponential function and obtained the best-fit parameter values: for the slope *κ* =0.00172 and the intercept *H* _0_=1816 mm^2^. The spatial block had a significant effect on *κ* (likelihood ratio 6.9, *p*=0.008) and *H* _0_ (likelihood ratio 41, *p*=1.4 × 10^*−*10^), but the cultivar had a much greater effect on *κ* (likelihood ratio 856, *p*=1.3 *×* 10^*−*47^) and *H* _0_ (likelihood ratio 2636, *p*<10^*−* 50^).

**Figure 3.**
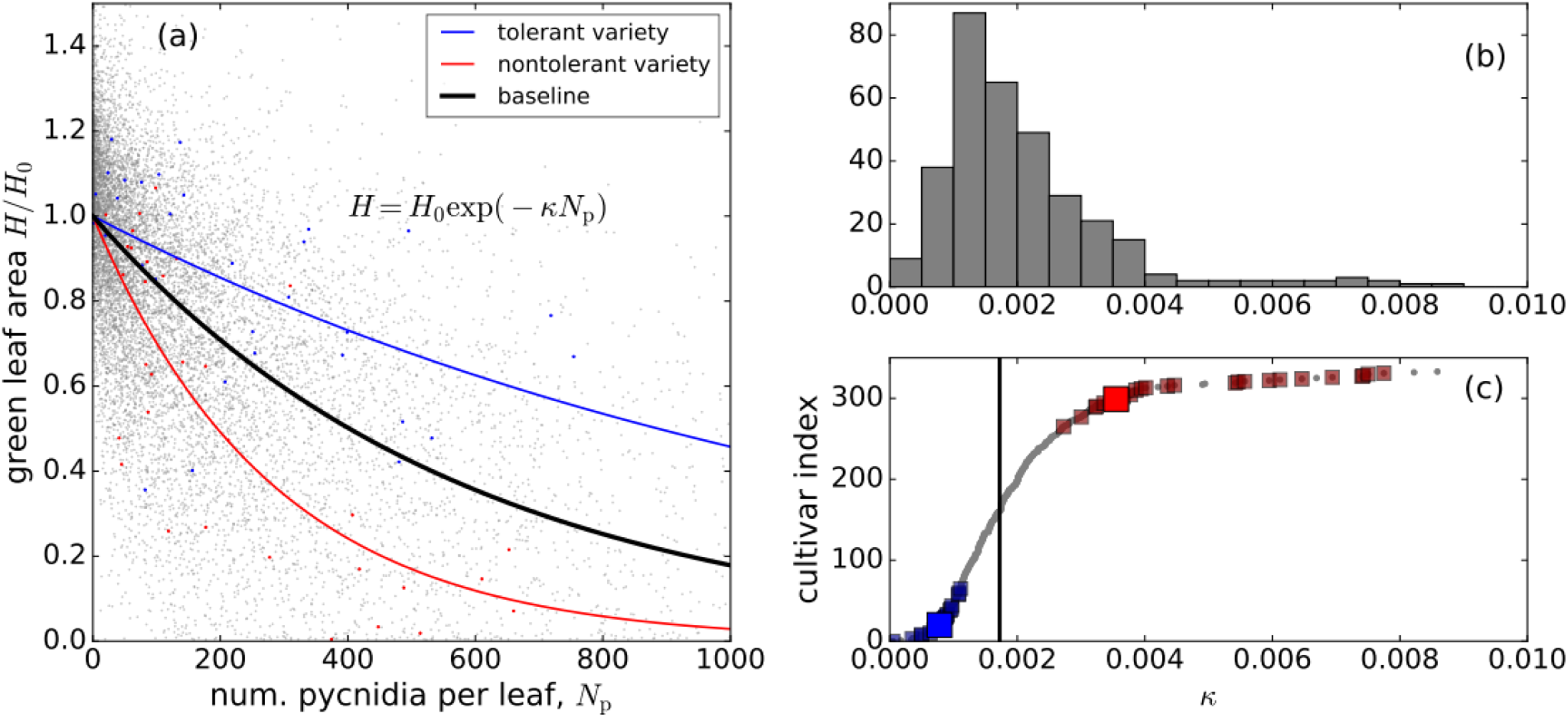
Tolerance of wheat to *Z. tritici* measured on the scale of individual leaves. (a) Green (Healthy) leaf area normalized by the leaf area in the absence of disease, *H*/ *H*_0_, is plotted versus the number of pycnidia per leaf *N*_*p*_. 11152 individual leaf measurements are shown using grey points. Best fit curves based on the exponential function *H* / *H* _0_=exp [− *κ N*_*p*_] are shown for all data (black curve) and for two example cultivars, a tolerant cultivar (Intact, blue curve), and an intolerant cultivar (Lynx, red curve). (b) The distribution of the 335 cultivars with respect to their *κ*-estimates. The most tolerant cultivars are those with the lowest *κ*-estimates at the left of the distribution. (c) Ranking of wheat cultivars according to their tolerance. *κ*-estimates for the 335 cultivars are shown in order of decreasing tolerance, that is increasing slope *κ* (grey points). Cultivars with tolerance significantly different from the baseline tolerance (black line) are marked with blue points (more tolerant) and red points (less tolerant), according to one-sided bootstrap t-tests with the confidence threshold of 0.05. Cultivars illustrated in panel (a) are marked using larger blue (cultivar Intact) and red (cultivar Lynx) squares.

There does not appear to be a clear pattern in terms of the goodness of fit neither for cultivars with different levels of tolerance nor for cultivars with different levels of resistance. To illustrate this, we present the goodness of fit metrics for both fit functions for four cultivars representing contrasting levels of tolerance and resistance. In a more tolerant cultivar Intact (blue curve in Fig. 3c), the linear fit yields *s*=349, *R*^2^=0.26, while the exponential fit yields *s*=345, *R*^2^=0.28. But in a less tolerant cultivar Lynx (red curve in Fig. 3c), the linear fit gives *s*=430, *R*^2^=0.59 compared to the exponential fit that gives *s*=415, *R*^2^=0.62. In a more resistant cultivar Element the linear fit gives *s*=450, *R*^2^=0.2 and the exponential fit gives *s*=448, *R*^2^=0.21; in a less resistant Arack the linear fit gives *s*=496, *R*^2^=0.37, while the exponential fit gives *s*=490, *R*^2^=0.4. The fits for these two cultivars are shown in Fig. S2.

### Ranking of cultivars according to their tolerance to *Z. tritici*

We estimated *κ* for each cultivar by fitting the empirical dependency of the green leaf area on the number of pycnidia with the exponential function [Eq. (3)], i.e. we obtained tolerance curves for each cultivar (such as the two tolerance curves depicted Fig. 3a in blue and red). The distribution of the *κ*-estimates is shown in Fig. 3b. Next, we ranked the cultivars according to their tolerance to *Z. tritici* (see Fig. 3c and Table S1). Smaller *κ*-values corresponded to more tolerant cultivars. We also compared *κ*-estimates for each cultivar to the baseline value (black vertical line in Fig. 3c). We found that 22 cultivars were significantly more tolerant than the baseline (blue squares in Fig. 3c) and 25 cultivars were significantly less tolerant than the baseline (red squares in Fig. 3c). Thus, the cultivars that we investigated in our field experiment exhibited significant differences with respect to leaf tolerance.

To determine to what extent the ranking of cultivars with respect to their *κ*-estimates was conserved between the two replicate blocks, we estimated the *κ*-values for each cultivar separately in each of the blocks. The *κ*-estimates exhibited a positive and significant correlation between the two replicates (*r*_*S*_ =0.18, *p*=0.001). In addition, we obtained similar data for a subset of 38 cultivars in 2015 (Stewart et al., 2016), which allowed us to evaluate the robustness of the outcomes. The *κ*-estimates exhibited a positive but a non-significant correlation between the two years (*r*_*S*_ =0.3, *p*=0.07).

### Relationship between tolerance/resistance and the year of cultivar registration

In a subset of 205 out of 335 cultivars, we had information on cultivar registration years. In those cultivars, tolerance increased with the year of cultivar registration: the correlation between the intolerance parameter *κ* and the cultivar’s registration year was negative and significant (r_S_ = −0.17, p = 0.02). In contrast, resistance did not exhibit a significant correlation with the cultivar’s registration year (r_S_ = −0.06, p = 0.36).

### Evidence for a tradeoff between leaf tolerance and resistance

We found that the estimates of tolerance, *κ*, for each of the 335 cultivars correlated negatively with the mean number of pycnidia per leaf, *N*_*p*_, the measure of resistance to STB, with *r*_*S*_ =*−* 0.27, *p*=5.8 *×* 10^*−*7^ (Fig. 4). Interestingly, *κ*-estimates correlated positively with mean PLACL values in each cultivar (*r*_*S*_ =0.31, *p*=8.6 *×* 10^*−*9^).

**Figure 4.**
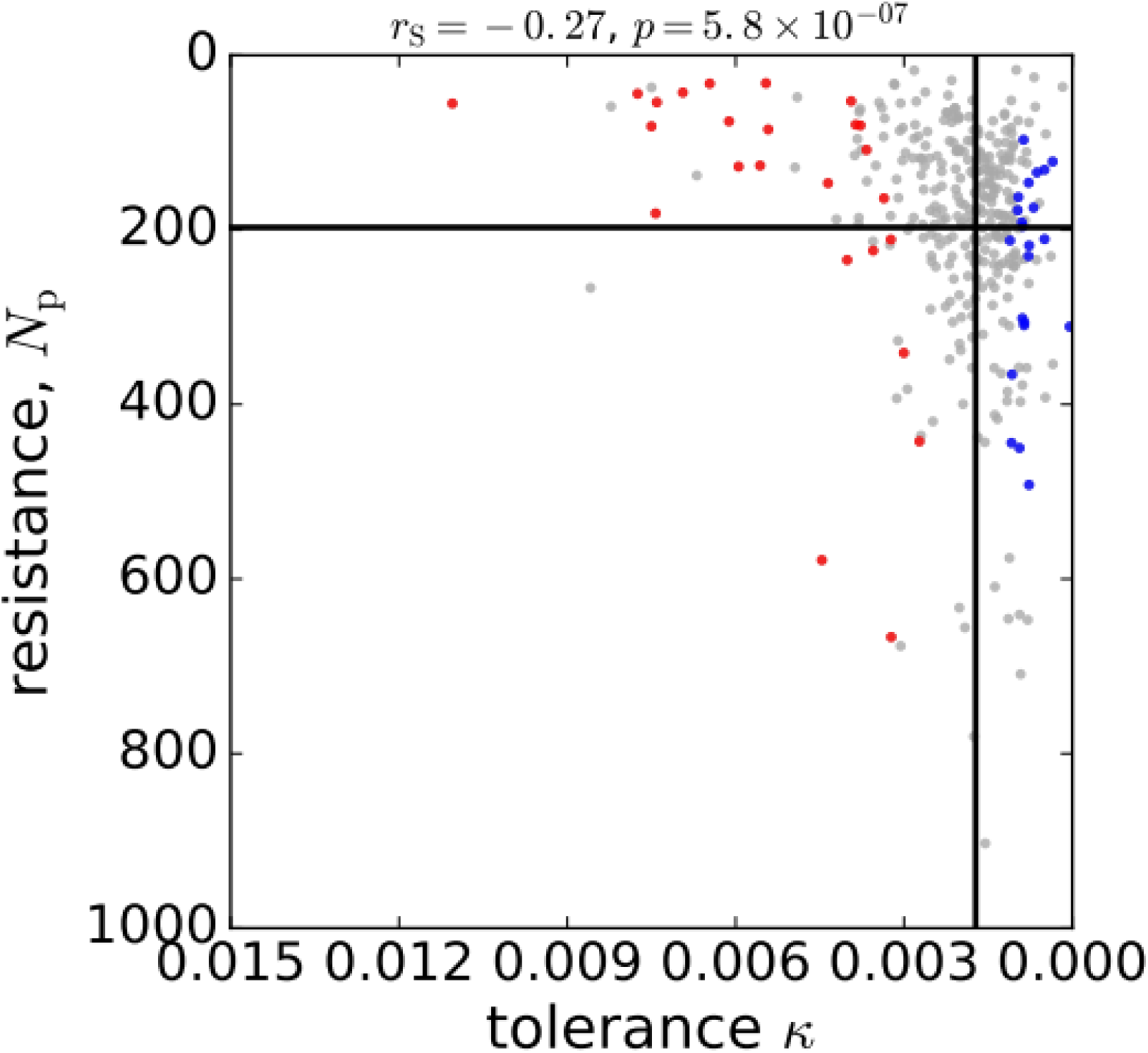
Correlation between leaf tolerance and resistance of wheat to STB. The measure of resistance, *N*_*p*_ is plotted against the measure of tolerance, *κ*, for each of 335 wheat cultivars (grey circles). Horizontal line shows the mean resistance and vertical line shows the baseline tolerance; blue/red circles mark the cultivars that are significantly more/less tolerant than the baseline.

Why are more tolerant cultivars on average less resistant? In other words, why do more tolerant cultivars carry more pycnidia on their leaves than less tolerant cultivars? A possible explanation is that since the leaf area is limited, this places a constraint on the maximum number of pycnidia that a leaf can carry. This constraint should lead to a negative relationship between *N*_*p*_ and *κ* in cases where the number of pycnidia on leaves belonging to the same cultivar approach the maximum allowed by the leaf area. A less tolerant cultivar suffers a larger necrotic area forming, on average, per pycnidium [parameter *a*_*p*_ in Eq. (1)]. Consequently, the maximum number of pycnidia per leaf is lower in a less tolerant cultivar than in a more tolerant cultivar. Therefore, the limitation in the leaf area should affect more strongly pathogen populations infecting less tolerant cultivars, where the green leaf area decreases more steeply with increasing numbers of pycnidia (Fig. 5a). If the limitation in the leaf area is indeed the dominant factor responsible for the negative relationship between tolerance and resistance, then the negative correlation should be present in intolerant cultivars, but absent in tolerant cultivars. This is because only intolerant cultivars have a high proportion of their leaf area covered by lesions already at rather modest numbers of pycnidia (red curves in Fig. 5a).

**Figure 5.**
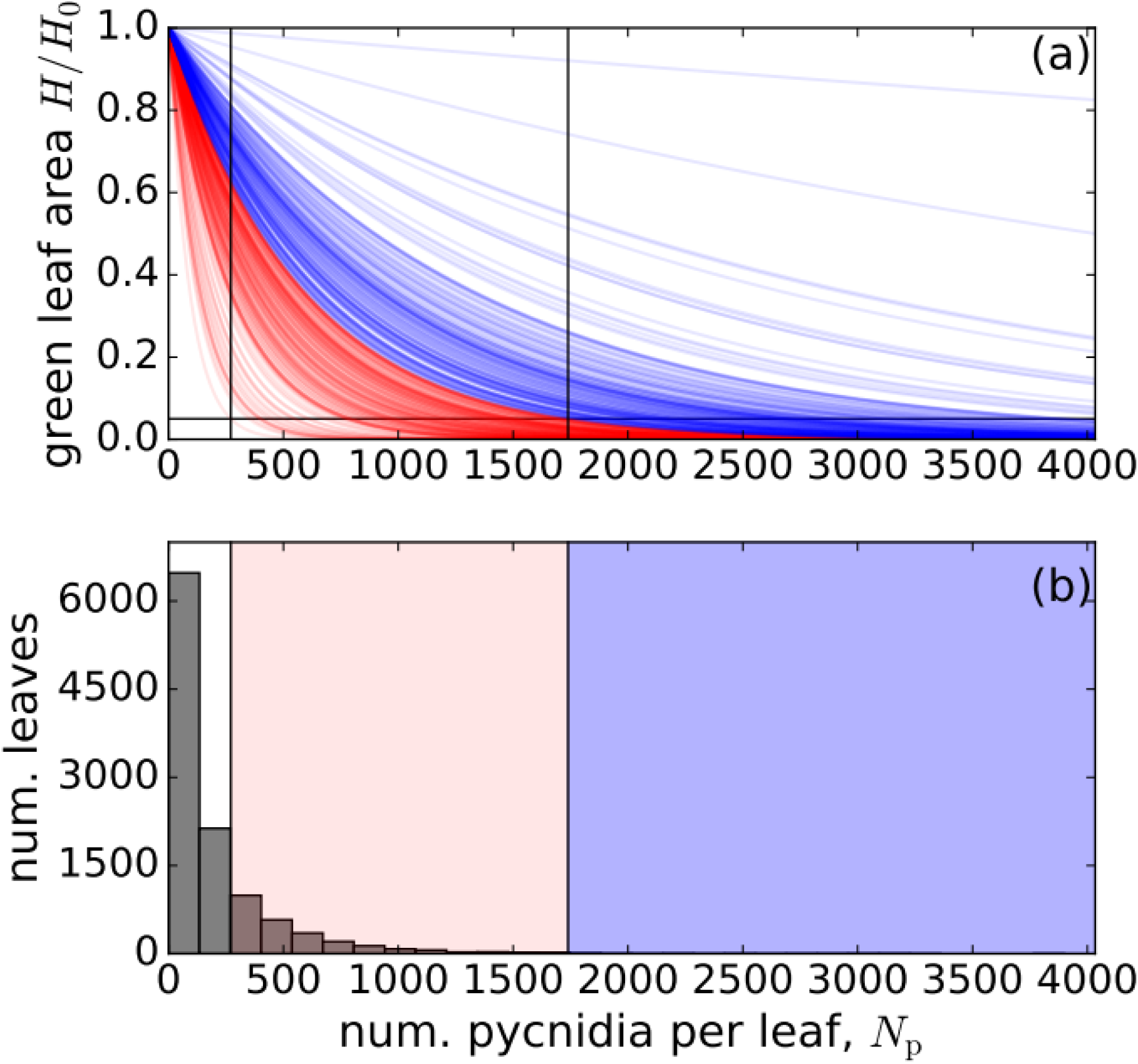
Limitation in the leaf area in tolerant and intolerant wheat cultivars. (a) Tolerance curves fitted to individual leaf data (raw data points are shown in Fig. 3a) for 335 wheat cultivars, blue curves correspond to cultivars more tolerant than the baseline (black curve) and red curves correspond to cultivars less tolerant than the baseline. The horizontal line shows the threshold *H*/*H*_0_=0.05. (b) Distribution of 11,152 individual infected leaves with respect to the number of pycnidia per leaf, *N*_p_. Shaded areas illustrate the ranges of *N*_p_-values in which the pathogen population infecting tolerant (blue) and intolerant (red) wheat varieties is affected by the limitation in the leaf area.

To test this expectation in a more quantitative fashion, we subdivided all cultivars into two groups according to their tolerance estimates using the baseline tolerance (*κ* =0.00172) as the threshold. Next, we conducted the Spearman’s correlation test (based on a t-test) in each of the groups separately. We found that intolerant cultivars exhibited a significant correlation between tolerance and resistance (*r*_*S*_ =*−* 0.34, *p*=4.3 *×* 10^*−*6^), while tolerant cultivars showed no significant correlation between the two traits (*r*_*S*_ =*−* 0.09, *p*=0.27). To test the validity of this outcome, we performed a series of more robust tests based on a bootstrap t-test. We first computed the uncertainty in the estimate *r*_*S*_ =*−* 0.27 for all cultivars in the form of the 95 % confidence interval: CI, −0.37 to −0.16. We also computed the uncertainties in *r*_*S*_ -estimates for the two groups of cultivars, tolerant (*r*_*S*_ =*−* 0.09, CI, −0.25 to 0.07) and intolerant (*r*_*S*_ =*−* 0.34, CI, −0.47 to −0.2). Figure S1 visualizes the bootstrap distributions of *r*_*S*_ and the CIs of *r*_*S*_ -estimates. The bootstrap t-test confirmed the outcome of the conventional t-test: the correlation between tolerance and resistance was significant among intolerant cultivars (*p*=4.0 *×*10^*−*6^) and not significant among tolerant cultivars (*p*=0.28). Furthermore, we used a more stringent bootstrap t-test to compare the *r*_*S*_ -estimates among tolerant and intolerant cultivars and found that the correlation among intolerant cultivars was significantly more negative than among tolerant cultivars (*p*=0.019).

Figure 5 illustrates why intolerant cultivars should be more strongly affected by the limitation in the leaf area than tolerant cultivars. Figure 5a shows the tolerance curves for tolerant (blue) and intolerant (red) cultivars. We determined the maximum number of pycnidia that can be reached in each cultivar by computing the number of pycnidia, *N*_pm_, at which 95 % of the green leaf area is lost on average (this is given by the intersection of each tolerance curve with the horizontal line *H*/*H*_0_=0.05 in Fig. 5a). Next, we computed the ranges in terms of *N*_pm_ corresponding to tolerant cultivars (blue-shaded area in Fig. 5b) and intolerant cultivars (red-shaded area in Fig. 5b). To determine the extent to which pathogen populations infecting tolerant and intolerant cultivars could be affected by the limitation in the leaf area, we compared these ranges with the overall distribution of leaves with respect to the numbers of pycnidia they carry (cf. the histogram in Fig. 5b and the blue- and red-shaded areas). While the blue-shaded area in Fig. 5b contained 38 leaves that constitute only about 0.3 % of the total number of 11,152 infected leaves, the red-shaded area contained a much more substantial proportion (22 %) of the leaves (2,485 leaves out 11,152). Therefore, the intolerant cultivars contained a much greater proportion of the leaves on which pathogen populations were likely to be affected by limitations in the leaf area compared to the tolerant cultivars. Combined with the observation that only intolerant cultivars exhibited a negative relationship between tolerance and resistance, these data support our hypothesis that the negative relationship between tolerance and resistance arises largely due to a limitation in the leaf area.

## Discussion

We discovered a novel component of tolerance in wheat to *Z. tritici* that operates on the scale of individual leaves (leaf tolerance). We devised an approach to quantify leaf tolerance empirically based on automated measurements of the green leaf area and the numbers of pycnidia on individual leaves. We gathered data from 11,152 individual infected leaves and characterized leaf tolerance in 335 elite European wheat cultivars. Cultivars exhibited significant differences in leaf tolerance, suggesting that this trait is at least partially under genetic control. We also found a negative relationship between leaf tolerance and resistance to *Z. tritici*, indicating that there is a tradeoff between tolerance and resistance. Our study presents the first clear evidence for such a tradeoff in the context of plant-pathogen interactions. We discuss the consequences of this possible tradeoff for the selection of tolerance/resistance in agricultural host populations.

Surprisingly, the nature of this tradeoff turned out to be different from what we expected based on ecological theory. Our analysis shows that the tradeoff is only present in cultivars with less than average tolerance (intolerant cultivars) and that the limitation in the leaf area is the dominant factor responsible for its occurrence. This mechanism differs from the host metabolic constraints that are usually implicated in the tradeoff between resistance and tolerance. We expect this novel mechanism underlying a tradeoff between tolerance and resistance could operate across a large class of infectious diseases in plants and animals, in which tolerance to the pathogen can be measured and the amount of host resources available to the pathogen can limit the pathogen population within the host. A key conceptual outcome of our study is that observing a negative relationship between tolerance and resistance is not necessarily indicative of a metabolic tradeoff, whereby tolerance and resistance confer fitness costs. Instead, as we show here, a tradeoff can arise via an entirely different mechanism, namely a limitation in the amount of host tissue or resources available to the pathogen (or more generally a limitation in the degree of fitness a host can lose because of infection). In the two prominent examples of a negative relationship between tolerance and resistance found in the literature (herbivory in plants, Fineblum & Rausher, 1995; malaria in mice, Råberg et al., 2007), the mechanisms underlying this relationship remain unknown.

The limitation in the leaf area is expected to constrain the evolution of pathogen populations towards higher reproductive fitness on intolerant cultivars, but the pathogen may overcome this limitation by evolving lower virulence (Anderson & May, 1982). According to our current understanding in ecological theory, a metabolic tradeoff between tolerance and resistance is expected that arises due to limitation in resources available to the host. Our data does not exclude the possibility of a metabolic tradeoff, but its detection may require an even more comprehensive dataset than what we have at hand. Evidence for the metabolic tradeoff is more likely to be found in the future by considering a larger number of sufficiently tolerant cultivars, because as we demonstrated here, in more tolerant cultivars the relationship between tolerance and resistance is not dominated by the limitation in the leaf area.

Jackson et al. (2014) reported that mature male voles were more tolerant to macroparasite infection compared to young males. Zeller & Koella (2017) found that the availability of nutrients influenced the magnitude of tolerance to microsporidian parasites in mosquito populations: mosquitos that had a restricted food supply were generally less tolerant to infection. It is plausible that both of these factors influence the tolerance of wheat leaves to *Z. tritici*. First, only the STB-induced damage on the three upper-most wheat leaves correlates strongly with yield loss (e.g., Thomas et al., 1989). Hence leaf tolerance may confer a fitness advantage to plants only during the later developmental stages when these leaf layers have already emerged. As a result, selection may have favored tolerance to manifest only during this late stage of development (similar to the adult plant resistance that is well known for several plant diseases). Second, the severity of STB epidemics is known to increase with increased rates of nitrogen fertilization (Leitch & Jenkins, 1995). This may result from an improved nutritional or physiological status of the leaves or a more disease-conducive physical environment. Hence leaf tolerance and its relationship with resistance may be affected by changing the rate of nitrogen application. Empirical investigation of both of these factors is feasible in the *Z. tritici*-wheat pathosystem and would improve our understanding of the ecological determinants of tolerance.

The dataset we used to characterize tolerance to a plant pathogen is unusually large compared to previous studies. For example, the number of different wheat genotypes used to study tolerance of wheat to STB in earlier studies ranged from 2 to 25 (Eyal & Ziv, 1974; Zuckerman et al., 1997; Parker et al., 2004; Foulkes et al., 2006; Collin et al., 2018). Råberg et al. (2007) investigated tolerance of mice to malaria infection using five mouse strains and three strains of *Plasmodium chabaudi*. Only the study of human tolerance to HIV (Regoes et al., 2014) and the study of tolerance in the wild population of Soay sheep to a gastrointestinal nematode infection (Hayward et al., 2014) had comparably large datasets that included thousands of infected individuals. Remarkably, as we demonstrated here, both tolerance and resistance can be readily quantified from digital images of infected leaves.

Our analyses and interpretations are based on two important assumptions: (i) the reduction in the green area of the second leaf is a major driver of yield loss induced by STB, and; (ii) the number of pycnidia per leaf is a good measure of the size of the pathogen population on a leaf. We justify these assumptions as follows: (i) Compared to infected leaves that have a large fraction of their surface area covered by lesions, leaves with a larger green area intercept a larger fraction of the incoming radiation, which contributes to plant yield. There is overwhelming empirical evidence showing that the reduction in the green leaf area is a major driver of yield loss for many leaf-affecting diseases of wheat (for example, Teng and Gaunt, 1980; Seck et al., 1991; Gaunt 1995; Bhathal et al., 2003), including STB (Eyal & Ziv, 1974; King et al., 1983; Forrer & Zadoks, 1983; Shaw & Royle, 1989b; Thomas et al., 1989). In particular, these studies conclude that the reduction in yield is strongest for the three upper leaves (including the second leaf on which we focused in this study) if the green leaf area is measured during the critical phase of seed development. In our field experiment, we could not determine yield corresponding to each individual infected leaf that we sampled. However, we measured overall yield per plot and found significant correlations between the green leaf area of second leaves (sampled at GS 75-85) and yield, measured both as tons per hectare and as thousand kernel weight (*r*_*S*_ =0.13, *p*=0.017 for yield measured in t/ha and *r*_*S*_ =0.13, *p*=0.019 for yield measured as TKW, see Results, Fig. 2). Note that the green leaf area was recorded only on infected leaves, while the yield was measured from plants sampled without regard to their infection status, hence the samples used to calculate yield comprised both healthy and infected plants. Therefore, the correlation coefficients we obtained here are likely to considerably underestimate the actual correlations between the green leaf area and yield, consistent with previous studies that found much stronger correlations (Eyal & Ziv, 1974; King et al., 1983; Forrer & Zadoks, 1983; Shaw & Royle 1989b). In particular, Thomas et al., (1989) reported that STB severity on second (flag-1) leaves had a particularly strong effect on yield. Thus, there is compelling empirical evidence in the existing literature and also an indication in the present study showing that the reduction in the green leaf area of second leaves contributes substantially to the yield loss induced by the disease.

To justify (ii) we first note that in this study we investigate tolerance and resistance from an evolutionary perspective. Hence, the measure of pathogen burden should reflect the reproductively active population of the pathogen. Measuring the total number of spores produced per leaf may provide a better way to quantify pathogen burden, but was not possible in our experiment because it could not be automated. However, we believe that the number of pycnidia is a reasonable proxy of pathogen burden because the number of pycnidia was shown to be the main factor determining the number of pathogen spores produced on an infected leaf (Stewart et al., 2016a). In addition, a recent field experiment showed that the proportion of the leaf area covered by STB lesions was largely independent from the number of pycnidia produced on a leaf (Karisto et al., 2018). Combining these two findings led us to conclude that the number of pycnidia per leaf is a better measure to quantify the pathogen population inhabiting a leaf than the area of a leaf damaged by infection.

According to our statistical analysis, an exponential decrease better fits the empirical dependency of the green leaf area on the number of pycnidia per leaf than a linear decrease, demonstrating that leaf tolerance curves were nonlinear. This deviates from what was reported in earlier analyses of wheat tolerance to *Z. tritici*: tolerance curves were typically fitted using linear functions (Eyal & Ziv, 1974; Parker et al., 2004; Foulkes et al., 2006), with the notable exception of the study by Shaw & Royle (1989b) that used a family of nonlinear curves. It was important to establish the departure from linearity in our study for two reasons. First, it allowed a more accurate comparison of tolerance estimates in different cultivars against the baseline. Second, it provided additional insight into the biology of the infection, because the linear model and the exponential model are based on different biological assumptions.

Our analysis of the model (Notes S1) demonstrates that the linear function Eq. (2) approximates well the relationship between the green leaf area and the number of pycnidia on the leaf when the number of pycnidia on the leaf is sufficiently low. This implies that the number of lesions on the leaf is also likely to be low, with the necrotic area covering only a small proportion of the total leaf area. Under this scenario, lesions develop mostly independently of each other. However, when lesions start to occupy a large proportion of the total leaf area, they become more likely to influence each other’s development due to limitations in space and/or resources. Under this scenario, Eq. (2) is no longer a good approximation and the necrotic area *a*_*p*_=*a*_*l*_/*n*_*p*_ that corresponds to a single pycnidium may depend on both green leaf area *H* and the number of pycnidia *N*_*p*_ already present on the leaf (i.e., a density dependence). Above, we considered the simplest case of this dependency when *a*_*p*_ is proportional to *H*, which resulted in the exponential solution [Eq. (3)]. This dependency may result from the lesion area *a*_*l*_ being proportional to the remaining green leaf area *H*. Biologically, this means that as more of the green leaf area becomes occupied by lesions, lesions tend to grow to a smaller size due to limitations in available green space and/or resources in the leaf. Alternatively, this dependency may arise due to the number of pycnidia per lesion *n*_*p*_ being inversely proportional to the remaining green leaf area *H*. This may occur due to an increased activation of plant defenses as more of the leaf area becomes occupied by lesions. Since our analysis shows that Eq. (3) is better supported by the data we collected for *Z. tritici* than Eq. (2), we conclude that density dependence contributes to the relationship between the number of pycnidia and the green leaf area, and is therefore expected to influence epidemiological dynamics on the scale of individual leaves. However, dedicated experiments under controlled conditions will be needed to reveal the mechanism behind the density dependence.

We recently identified several chromosomal regions and candidate genes in the wheat genome associated with resistance to STB (quantified as the mean pathogen burden, i.e. the mean number of pycnidia per leaf) using the same phenotypic dataset (Karisto et al., 2018) and a genome-wide association study (GWAS; Yates, et al., 2019). We hypothesize that a GWAS based on the leaf-level tolerance estimates that we report here could also identify significantly associated chromosomal regions. This would indicate that leaf-level tolerance has an underlying genetic basis and is subject to evolutionary processes, and potentially elucidate molecular mechanisms affecting leaf-level tolerance. One possible mechanism could be related to additive actions of toxin sensitivity genes carried by different wheat cultivars that interact with host-specific toxins produced by the pathogen, as demonstrated for *Parastagonospora nodorum* on wheat (Friesen et al., 2008; Oliver et al., 2012). This mechanism would contribute to tolerance if the number of actively interacting toxin - toxin sensitivity gene pairs exceeds a threshold beyond which the removal of a single gene pair does not impair the pathogen reproduction, but nevertheless reduces the host damage, thereby decreasing the average necrotic area per pycnidium and increasing leaf tolerance.

Breeding for resistance to STB disease is based on disease assessments that do not quantify pathogen reproduction on the leaves. The amount of disease is typically assessed visually using a categorical scale of severities corresponding to different ranges in terms of the proportion of necrotic area on the leaves (or PLACL). As a result, breeders are likely to select for cultivars with lower PLACL. But as we have shown above, cultivars with lower PLACL are on average more tolerant. Hence, by focusing on PLACL wheat breeders may have inadvertently selected for increased tolerance. Due to the tradeoff between tolerance and resistance, this simultaneously favors lower levels of STB resistance. Some support for this hypothesis is given by our preliminary analysis of the relationship between tolerance/resistance and the year of cultivar registration. In a subset of 205 out of 335 cultivars, we found that estimates of tolerance increased with the year of cultivar registration, while estimates of resistance did not exhibit a significant change over time *κ*. This pattern suggests that the selection practices used by plant breeders have led to wheat populations in Europe with higher tolerance but lower resistance to STB over time.

The method to quantify leaf tolerance presented here can potentially be used to measure tolerance to other pathogens that infect plant leaves. The necessary condition is that digital images of infected leaves should enable quantitative measurements of both the damage to the plant induced by the pathogen and the size of the pathogen population on the leaf. This should be possible for many necrotrophic pathogens that form visible fruiting bodies on the leaf surface. Using this approach may facilitate the discovery of similar tradeoffs between tolerance and resistance across a wide array of plant-pathogen systems.

## Supporting information

Table S1. Ranking of wheat cultivars according to their tolerance to Zymoseptoria tritici.

## Acknowledgements

We thank P. Karisto, D. Dos Santos Pereira, S. Fouché, and L. Meile for help in collecting and processing leaf samples; H. Zellweger for managing the wheat trial; and A. Hund and J. Anderegg for providing data on wheat yield. A. Mikaberidze would like to thank P. Karisto and R. Regoes for fruitful discussions. A. Mikaberidze gratefully acknowledges financial support from the Swiss National Science Foundation through the Ambizione grant PZ00P3_161453.

## Author Contributions

AM and BAM conceived and designed the experiment. AM supervised the field collection and processing of leaf samples. AM analyzed the data in discussions with BAM. AM wrote the manuscript. AM and BAM revised the manuscript.

## Data Accessibility Statement

Raw data is available from the Dryad Digital Repository: https://doi.org/10.5061/dryad.171q4. We confirm that, should the manuscript be accepted, additional data supporting the results will be archived in an appropriate public repository such as Dryad or Figshare, and the data DOI will be provided at the end of the article.

## Supporting Information

The following Supporting Information is available for this article:

**Notes S1.** Simple model of leaf tolerance

**Notes S2.** Two components of tolerance

Figure S1. Detailed statistical analysis of the correlation between tolerance and resistance among tolerant and intolerant cultivars.

Figure S2. Example fits for cultivars with contrasting levels of resistance.

**Table S1.** Ranking of wheat cultivars according to their tolerance to *Zymoseptoria tritici*.

**Notes S1.** Simple model of leaf tolerance

Here we derive a simple model of infection of wheat leaves by *Z. tritici*. We assume that each lesion has an area *a*_*l*_ and contains *n*_*p*_ pycnidia. How much will the green leaf area *H* diminish under a small increase in the number of pycnidia *Δ N*_*p*_? To find out, we express a small decrease in the green leaf area *ΔH* in terms of a small increase in the number of lesions: *ΔH* =*−a*_*l*_ *Δ N*_*l*_. At the same time, the increment in the number of pycnidia *Δ N*_*p*_ is related to the increment in the number of lesions: *Δ N*_*p*_ =*n*_*p*_ *Δ N*_*l*_. A simple rearrangement yields the relationship between the increment in the number of pycnidia and the decrease in the green leaf area: *ΔH* =*−a*_*l*_/*n*_*p*_ *Δ N*_*p*_. By taking the limit *Δ N*_*p*_ → 0 and *ΔH* → 0 we obtain the differential equation:

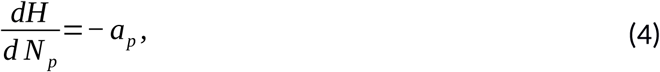

where *a*_*p*_=*a*_*l*_/*n*_*p*_ represents the area of the lesion that corresponds to a single pycnidium.

### Assumption 1

We first assume that *a*_*p*_ depends neither on *H*, nor on *N*_*p*_. In this case, the solution of Eq. (4) is a linear function that can be written as:

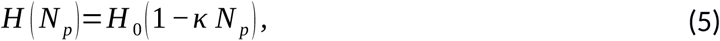

where *κ* =*a*_*p*_*H*_0_ and *H*_0_ is the green leaf area in the absence of infection.

### Assumption 2

Here, we relax Assumption 1 and consider the case when the area *a*_*p*_ is proportional to the green leaf area *H*, i. e.

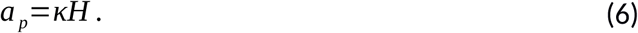

In this case, the solution of Eq. (4) is an exponential function that can be expressed as:

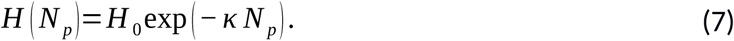

Note, that when *κ N*_*p*_ is small, we can expand the function *H* (*N*_*p*_) Eq. (7) in the Taylor’s series with respect to its argument *κ N*_*p*_. Retaining the zeroth and the first terms of the series yields the linear dependency Eq. (5).

**Notes S2**. Two components of tolerance

Here we demonstrate mathematically that the novel component of tolerance of wheat to STB that we measured in this experiment and the tolerance components that were measured previously contribute to overall tolerance as multiplicative factors. We use the critical point model (King et al.,1983; Shaw & Royle, 1989a) to relate the grain yield to the loss in the green leaf area in the three upper-most leaves:

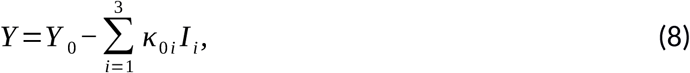

where *Y* is the yield in the presence of disease (measured for example in tons per hectare), *Y*_0_ is the yield in the absence of disease and *I*_*i*_ is the area of the *i*th leaf (i=1 for flag leaf, i=2 for flag-1 leaf, i=3 for flag-2 leaf) that became chlorotic or necrotic as a result of infection, measured during the critical stage of grain development (around GS 75). The slope *κ*_0*i*_ represents the intolerance parameter that may have different values in different leaf layers. We call this component of tolerance “whole-plant tolerance”, because it includes various mechanisms of compensation for the leaf damage at the level of the whole plant.

In this study, we argue that the number of pycnidia per leaf, *N*_*pi*_, corresponding to the *i*th leaf layer allows for a more accurate quantification of the pathogen population than the area of the leaf *I*_*i*_ damaged due to disease. Similarly to Eq. (8), this leads to the following relationship between the yield and the number of pycnidia per leaf:

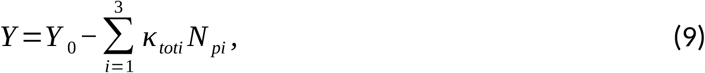

where *κ*_*toti*_ is the total intolerance parameter and as in Eq. (8), we perform a summation over the three upper-most leaf layers. Further, we assume for simplicity that the linear model Eq. (2) describes well the relationship between the green leaf area, *H*_*i*_, and the number of pycnidia per leaf *N*_*pi*_. The damaged leaf area, *I*_*i*_, can then be determined from Eq. (2), but for each individual leaf layer *i*, according to

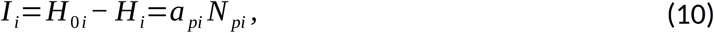

where *a*_*pi*_ is the damaged leaf area that corresponds on average to a single pycnidium, which quantifies the degree of tolerance on the scale of individual leaves that we measured in this study for the second leaf (i.e., flag-1 leaf). Here we used the linear approximation of the dependency of the green leaf area on the number of pycnidia, which works well when the number of pycnidia per leaf is sufficiently low. In this case, *a*_*pi*_ is related to the intolerance parameter, *κ*_*i*_, that we measured here through *H*_0*i*_, the green leaf area in the absence of disease: *a*_*pi*_=*H*_0*i*_*κ*_*i*_. To establish the relationship between the overall tolerance quantified by *κ_tot_*, whole-plant tolerance quantified by *κ*_0_ and leaf tolerance quantified by *κ*, we substitute *I* from Eq. (10) in Eq. (8)

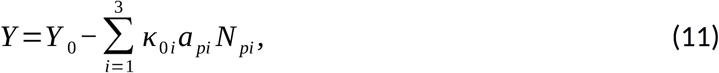

Comparison of Eq. (11) and Eq. (9) reveals that for each leaf layer *i*

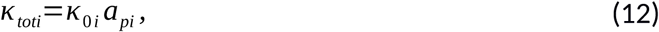

the total intolerance parameter is a product of the whole-plant and leaf intolerance parameters. If the relationships between the yield, *Y*, and the damaged leaf area, *I*_*i*_, and between the yield, Y, and the number of pycnidia per leaf, *N*_*pi*_, are not linear, then the yield would still be a decreasing, but a nonlinear function of the number of pycnidia per leaf, and the multiplicative relation between the total tolerance and the two components of intolerance in Eq. (12) will not be retained.

**Figure S1.**
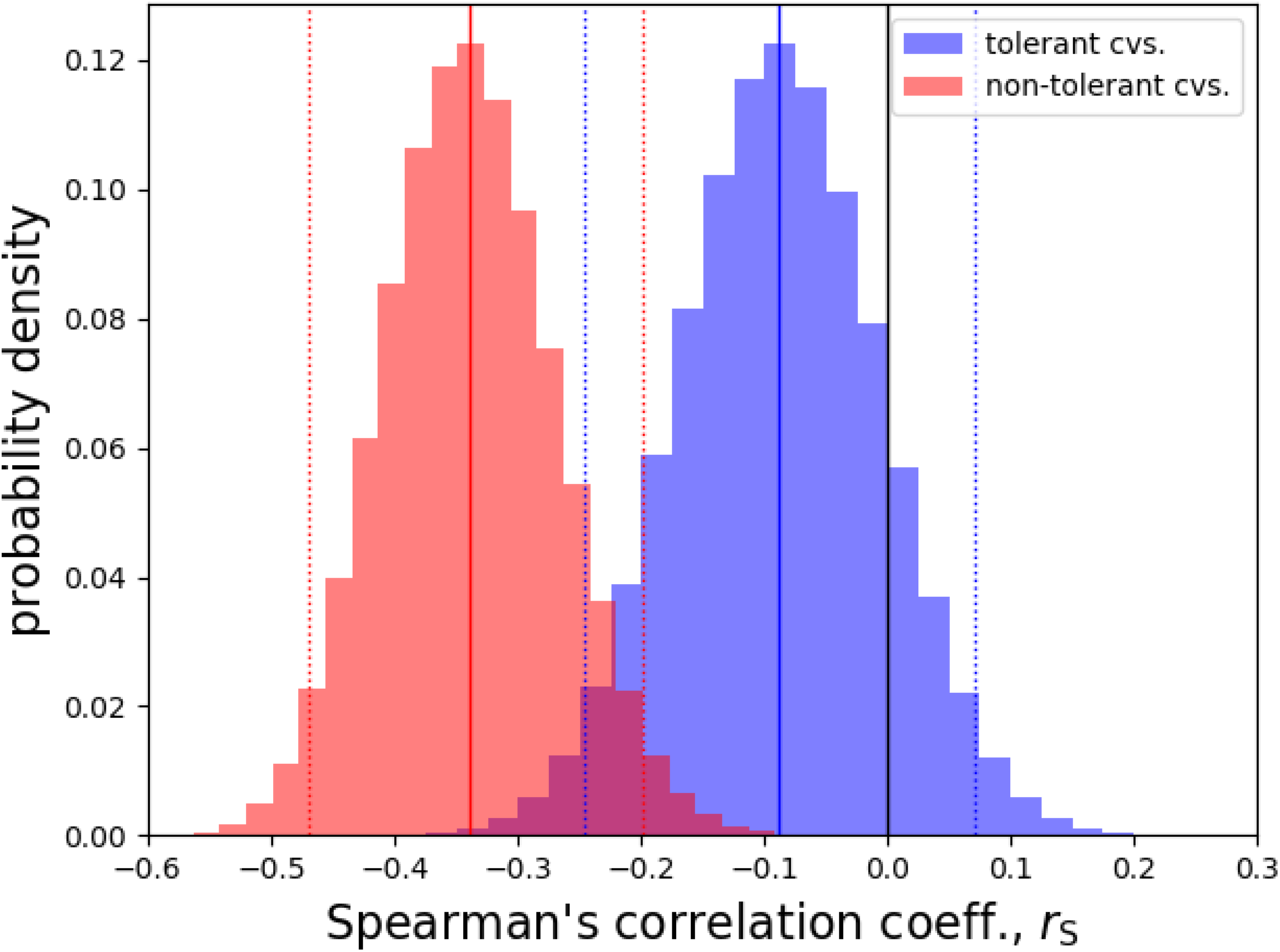
Detailed statistical analysis of the correlation between tolerance and resistance among tolerant and intolerant cultivars. The analysis is based on creating a large number of bootstrap samples (*n*_bs_=1’000’000) on the basis of the estimates of tolerance and resistance in 335 wheat cultivars shown in Fig. 3 of the main text. The histogram shows the distributions of the values of the Spearman’s correlation coefficient, *r*_S_, among tolerant (blue) and intolerant (red) cultivars. These distributions approximate the underlying probability distributions. Solid vertical lines show the point-estimates of *r*_S_ among tolerant (blue) and intolerant (red) cultivars. Dotted vertical lines show the 95 % confidence intervals of the corresponding point estimates.

**Figure S2.**
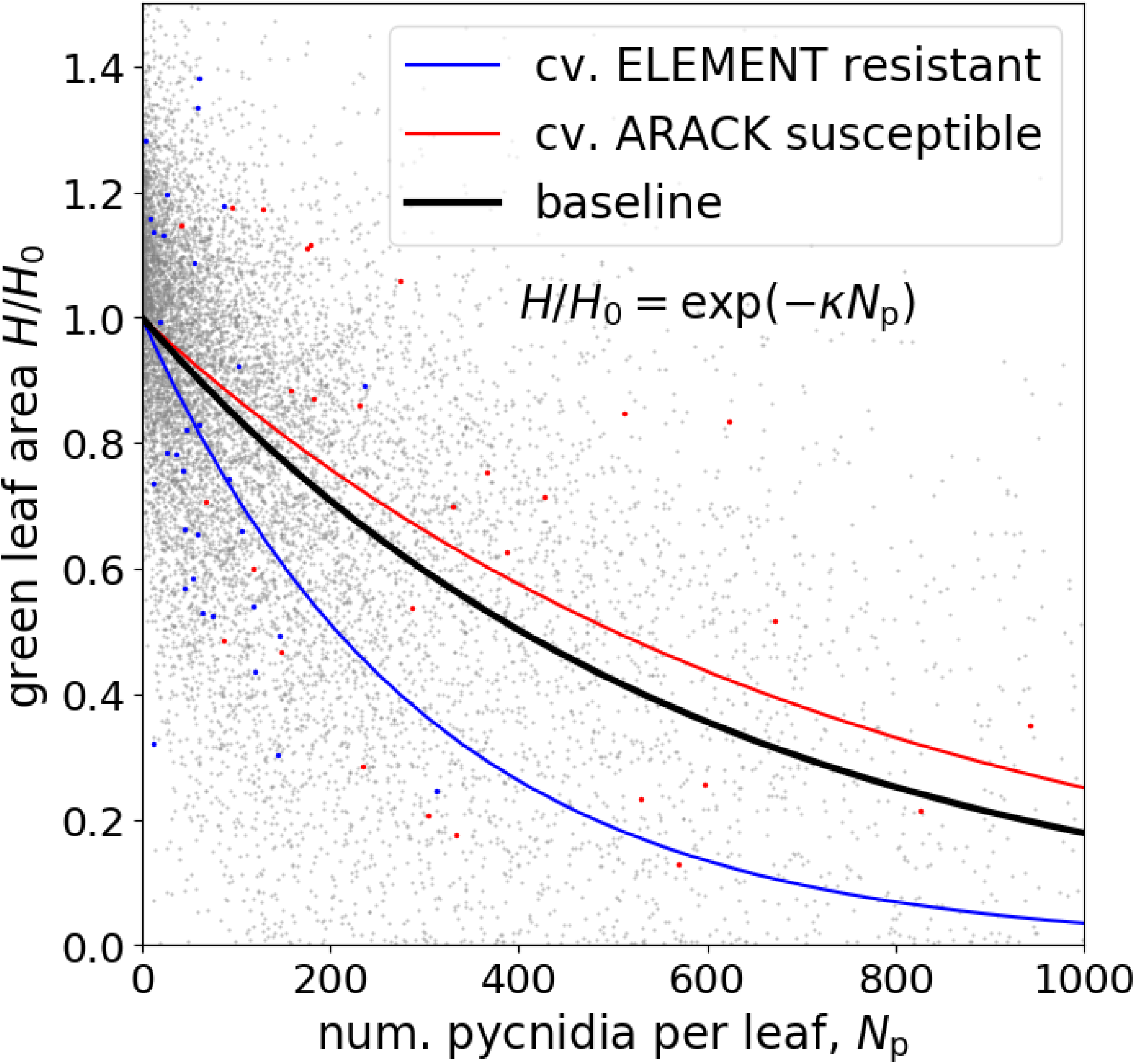
Example fits for cultivars with contrasting levels of resistance. Cultivar Element (estimate of resistance *N*_*p*_=66 estimate of tolerance *κ* =0.0033,) is more resistant than cultivar Arack (estimate of resistance *N*_*p*_=412, estimate of tolerance *κ* =0.0014). However, in terms of tolerance the two cultivars are not significantly different from the baseline (black curve).

## References

Anderson, R. M. & May, R.M. (1982). Coevolution of hosts and parasites. Parasitology, 85, 411–426.

Ayres, J.S. & Schneider, D.S. (2008). A Signaling Protease Required for Melanization in Drosophila Affects Resistance and Tolerance of Infections. PLoS Biol., 6, e305.

Ayres, J.S. & Schneider, D.S. (2012). Tolerance of Infections. Annu. Rev. Immunol., 30, 271–294.

Baucom, R.S. & De Roode, J.C. (2011). Ecological immunology and tolerance in plants and animals. Funct. Ecol., 25, 18–28.

Baucom, R.S. & Mauricio, R. (2008). Constraints on the evolution of tolerance to herbicide in the common morning glory: Resistance and tolerance are mutually exclusive. Evolution, 62, 2842–2854.

van den Berg, F, Paveley ND, Bingham IJ & van den Bosch F. (2017). Physiological Traits Determining Yield Tolerance of Wheat to Foliar Diseases. Phytopathology, 107, 1468–1478.

Best, A., White, A. & Boots, M. (2008). Maintenance of host variation in tolerance to pathogens and parasites. Proc. Natl. Acad. Sci., 105, 20786–20791.

Bhathal, J.S., Loughman, R. & Speijers, J. (2003). Yield reduction in wheat in relation to leaf disease from yellow (tan) spot and septoria nodorum blotch. European Journal of Plant Pathology, 109, 435–443.

Bingham, I.J., Walters, D.R., Foulkes, M.J. & Paveley, N.D. (2009). Crop traits and the tolerance of wheat and barley to foliar disease. Ann. Appl. Biol., 154, 159–173.

Blanchet, S., Rey, O. & Loot, G. (2010). Evidence for host variation in parasite tolerance in a wild fish population. Evol. Ecol., 24, 1129–1139.

Brown, J.K. (2002). Yield penalties of disease resistance in crops. Curr. Opin. Plant Biol., 5, 339–344.

Caldwell, R.M., Schafer, J.F., Compton, L.E. & Patterson, F.L. (1958). Tolerance to Cereal Leaf Rusts. Science, 128, 714–715.

Carr, D.E., Murphy, J.F. & Eubanks, M.D. (2006). Genetic variation and covariation for resistance and tolerance to Cucumber mosaic virus in *Mimulus guttatus* (Phrymaceae): A test for costs and constraints. Heredity, 96, 29–38.

Cobb, N.A. (1894). Contributions to an economic knowledge of Australian rust (Uredinae) Chap. 10. Agr. Gaz. N. S. W., 5, 239–50.

Collin, F., Bancal, P., Spink, J., Appelgren, P.K., Smith, J., Paveley, N.D., Bancal, M.O. & Foulkes, M.J. (2018). Wheat lines exhibiting variation in tolerance of Septoria tritici blotch differentiated by grain source limitation. Field Crop. Res., 217, 1–10.

Cowger, C., Hoffer, M.E. & Mundt, C.C. (2000). Specific adaptation by *Mycosphaerella graminicola* to a resistant wheat cultivar. Plant Pathol., 49, 445–451.

Davison, A. C. & Hinkley, D. V. (1997). Bootstrap methods and their applications. Cambridge University Press, Cambridge, UK.

Eyal, Z. & Ziv, O. (1974). The relationship between epidemics of Septoria leaf blotch and yield losses in spring wheat. Phytopathology, 64, 1385–1389.

Fineblum, W.L. & Rausher, M.D. (1995). Tradeoff between resistance and tolerance to herbivore damage in a morning glory. Nature, 377, 517–520.

Fones, H. & Gurr, S. (2015). The impact of Septoria tritici Blotch disease on wheat: An EU perspective. Fungal Genet. Biol., 79, 3–7.

Fornoni, J., Nunez-Farfan, J., Valverde, P.L. & Rausher, M.D. (2004). Evolution of Mixed Strategies of Plant Defense Allocation against Natural Enemies. Evolution, 58, 1685–1695.

Forrer, H. R. & Zadoks, J. C. (1983). Yield reduction in wheat in relation to leaf necrosis caused by Septoria tritici. Neth. J. Pl. Path., 89, 87–98.

Foulkes, M.J., Paveley, N.D., Worland, A., Welham, S.J., Thomas, J. & Snape, J.W. (2006). Major Genetic Changes in Wheat with Potential to Affect Disease Tolerance. Phytopathology, 96, 680–688.

Friesen, T. L., Faris, J. D., Solomon, P. S., and Oliver, R. P. (2008). Host-specific toxins: Effectors of necrotrophic pathogenicity, Cell. Microbiol., 10, 1421–1428.

Fraaije, B., Cools, H.J., Fountaine, J., Lovell, D.J., Motteram, J., West, J.S. & Lucas, J. A. (2005). Role of Ascospores in Further Spread of QoI-Resistant Cytochrome b Alleles (G143A) in Field Populations of *Mycosphaerella graminicola*. Phytopathology, 95, 933–941.

Gaunt, R. (1981). Disease Tolerance - an Indicator of Thresholds? Phytopathology, 71, 915–916.

Gaunt, R.E. (1995). The Relationship Between Plant Disease Severity and Yield. Annual Review of Phytopathology, 33, 119–144.

Hayward, A.D., Nussey, D.H., Wilson, A.J., Berenos, C., Pilkington, J.G., Watt, K.A., Pemberton, J.M. & Graham, A.L. (2014). Natural Selection on Individual Variation in Tolerance of Gastrointestinal Nematode Infection. PLoS Biol., 12, 1–13.

Herms, D.A. & Mattson, W.J. (1992). The Dilemma of Plants: To Grow or Defend. Q. Rev. Biol., 67, 283–335.

Howick, V.M. & Lazzaro, B.P. (2014). Genotype and diet shape resistance and tolerance across distinct phases of bacterial infection. BMC Evol. Biol., 14, 1–13.

Inglese, S.J. & Paul, N.D. (2006). Tolerance of *Senecio vulgaris* to Infection and Disease Caused by Native and Alien Rust Fungi. Phytopathology, 96, 718–726.

Jackson, J., Hall, A., Friberg, I., Ralli, C., Lowe, A., Zawadzka, M., Turner, A.K., Stewart, A., Birtles, R.J., Paterson, S., Bradley, J.E. & Begon, M. (2014). An immunological marker of tolerance to infection in wild rodents. PLoS Biol., 12, e1001901.

Jones E., Oliphant E., Peterson P. et al. (2001). SciPy: Open Source Scientific Tools for Python. Available at http://www.scipy.org/. Last accessed 30.05.2018.

Karisto, P., Hund, A., Yu, K., Anderegg, J., Walter, A., Mascher, F., McDonald, B.A. & Mikaberidze, A. (2018). Ranking quantitative resistance to Septoria tritici blotch in elite wheat cultivars using automated image analysis. Phytopathology, 108, 568–581

Kema, G.H., Yu, D., Rijkenberg, F.H., Shaw, M.W. & Baayen, R.P. (1996). Histology of the pathogenesis of mycosphaerella graminicola in wheat. Phytopathology, 86, 777–786.

King, J.E., Jenkins, J.E. & Morgan, W.A. (1983). The estimation of yield losses in wheat from severity of infection by Septoria species. Plant Pathology, 32, 239–249.

Kollers, S., Rodemann, B., Ling, J., Korzun, V., Ebmeyer, E., Argillier, O., Hinze, M., Plieske, J., Kulosa, D., Ganal, M.W. & Roder, M.S. (2013a). Genetic architecture of resistance to Septoria tritici blotch (*Mycosphaerella graminicola*) in European winter wheat. Mol. Breed., 32, 411–423.

Kollers, S., Rodemann, B., Ling, J., Korzun, V., Ebmeyer, E., Argillier, O., Hinze, M., Plieske, J., Kulosa, D., Ganal, M.W. & Roder, M.S. (2013b). Whole Genome Association Mapping of Fusarium Head Blight Resistance in European Winter Wheat (*Triticum aestivum L.*). PLoS One, 8.

Kover, P.X. & Schaal, B.A. (2002). Genetic variation for disease resistance and tolerance among *Arabidopsis thaliana* accessions. Proc. Natl. Acad. Sci., 199, 11270–11274.

Leitch, M.H. & Jenkins P.D. (1995). Influence of nitrogen on the development of Septoria epidemics in winter wheat. Journal of Agricultural Science, 124, 361–368.

Linde, C.C., Zhan, J. & McDonald, B.A. (2002). Population Structure of *Mycosphaerella graminicola*: From Lesions to Continents. Phytopathology, 92, 946–55.

Mauricio, R., Rausher, M.D. & Burdick, D.S. (1997). Variation in the defense strategies of plants: Are resistance and tolerance mutually exclusive?. Ecology, 78, 1301–1311.

Maze-Guilmo, E., Loot, G., Paez, D.J., Lefevre, T. & Blanchet, S. (2014). Heritable variation in host tolerance and resistance inferred from a wild host-parasite system. Proc. R. Soc. B Biol. Sci., 281, 20132567–20132567.

Medzhitov, R., Schneider, D.S. & Soares, M.P. (2012). Disease Tolerance as a Defense Strategy. Science, 335, 936–942.

van der Meijden, E., Wijn, M. & Verkaar, H.J. (1988). Defence and Regrowth, Alternative Plant Strategies in the Struggle against Herbivores. Oikos, 51, 3550–363.

Miller, M.R., White, A. & Boots, M. (2005). The evolution of host resistance: Tolerance and control as distinct strategies. J. Theor. Biol., 236, 198–207.

Newville, M., Stensitzki, T., Allen, D. B., & Ingargiola, A. (2014, September 21). LMFIT: Non-Linear Least-Square Minimization and Curve-Fitting for Python¶. Zenodo. http://doi.org/10.5281/zenodo.11813

Newton, A.C. (2016). Exploitation of Diversity within Crops: the Key to Disease Tolerance?. Front. Plant Sci., 7, 1–12.

Newton A.C., Guy D.C., Gaunt R.E., Thomas W.T.B. (2000). The effect of powdery mildew inoculum pressure and fertiliser level on disease tolerance in spring barley. Journal of Plant Diseases and Protection, 107, 67–73.

Ney, B., Bancal, M.O., Bancal, P., Bingham, I.J., Foulkes, J., Gouache, D., Paveley, N. & Smith, J. (2013). Crop architecture and crop tolerance to fungal diseases and insect herbivory. Mechanisms to limit crop losses. Eur. J. Plant Pathol., 135, 561–580.

Oliver, R. P., Friesen, T. L., Faris, J. D., and Solomon, P. S. (2012). *Stagonospora nodorum*: From pathology to genomics and host resistance. Annu. Rev. Phytopathol., 50, 23–43.

Pagan, I. & Garcia-Arenal, F. (2018). Tolerance to Plant Pathogens: Theory and Experimental Evidence. Int. J. Mol. Sci., 19, 810.

Pagan, I., Alonso-Blanco, C. & Garcia-Arenal, F. (2008). Host responses in life-history traits and tolerance to virus infection in *Arabidopsis thaliana*. PLoS Pathog., 4.

Parker, S.R., Shaw, M.W. & Royle, D.J. (1995). The reliability of visual estimates of disease severity on cereal leaves. Plant Pathol., 44, 856–864.

Parker, S.R., Welham, S., Paveley, N.D., Foulkes, J. & Scott, R.K. (2004). Tolerance of septoria leaf blotch in winter wheat. Plant Pathol., 53, 1–10.

R Core Team (2016). R: A Language and Environment for Statistical Computing. R Foundation for Statistical Computing, Vienna, Austria.

Råberg, L. (2014). How to Live with the Enemy: Understanding Tolerance to Parasites. PLoS Biol., 12.

Råberg, L., Sim, D. & Read, A.F. (2007). Disentangling Genetic Variation for Resistance and Tolerance to Infectious Diseases in Animals. Science, 318, 812–815.

Regoes, R.R., McLaren, P.J., Battegay, M., Bernasconi, E., Calmy, A., Gunthard, H.F., Hoffmann, M., Rauch, A., Telenti, A. & Fellay, J. (2014). Disentangling Human Tolerance and Resistance Against HIV. PLoS Biol., 12.

Restif, O. & Koella, J.C. (2004). Concurrent Evolution of Resistance and Tolerance to Pathogens. Am. Nat., 164, E90–E102.

Richardson, C.E., Kooistra, T. & Kim, D.H. (2010). An essential role for XBP-1 in hos protection against immune activation in *C. elegans*. Nature, 463, 1092–1095.

Roy, B.A., Kirchner, J.W., (2000). Evolutionary dynamics of pathogen resistance and tolerance. Evolution, 54, 51–63.

Roy, B.A., Kirchner, J.W., Christian C.E. & Rose, L.E. (2000). High disease incidence and apparent disease tolerance in a North American Great Basin plant community. Evolutionary Ecology, 14: 421–438

Schafer, J.F. (1971). Tolerance to Plant Disease. Annu. Rev. Phytopathol., 9, 235–252.

Schindelin, J., Rueden, C.T., Hiner, M.C. & Eliceiri, K.W. (2015). The ImageJ Ecosystem: An Open Platform for Biomedical Image Analysis. Mol. Reprod. Dev., 529, 518–529.

Seck, M., Roelfs, A.P. & Teng, P.S. (1991). Influence of leaf position on yield loss caused by wheat leaf rust in single tillers. Crop Protection, 10, 222–228.

Shaw, M.W. & Royle, D. (1989a). Airborne inoculum as a major source of Septoria tritici (*Mycosphaerella graminicola*) infections in winter wheat crops in the UK. Plant Pathology, 38, 35–43.

Shaw, M.W. & Royle, D.J. (1989b). Estimation and validation of a function describing the rate at which *Mycosphaerella graminicola* causes yield loss in winter wheat. Ann. Appl. Biol., 115, 425–442.

Shinzawa, N., Nelson, B., Aonuma, H., Okado, K., Fukumoto, S., Miura, M. & Kanuka, H. (2009). p38 MAPK-Dependent Phagocytic Encapsulation Confers Infection Tolerance in Drosophila. Cell Host Microbe, 6, 244–252.

Shuckla, A., Pagán, I. & García-Arenal, F. (2017). Effective tolerance based on resource reallocation is a virus-specific defence in *Arabidopsis thaliana*. Mol. Plant Pathol., 19, 1454–1465

Simms, E. & Triplett, J. (1994). Costs and Benefits of Plant Responses to Disease: Resistance and Tolerance. Evolution, 48, 1973–1985.

Soares, M.P., Teixeira, L. & Moita, L.F. (2017). Disease tolerance and immunity in host protection against infection. Nat. Rev. Immunol., 17, 83–96.

Stewart, E.L., Croll, D., Lendenmann, M.H., Sanchez-vallet, A., Hartmann, F.E., Palma-guerrero, J., Ma, X. & McDonald, B.A. (2016a). QTL mapping reveals complex genetic architecture of quantitative virulence in the wheat pathogen *Zymoseptoria tritici*. Mol. Plant Pathol., 19, 201–216.

Stewart, E.L., Hagerty, C.H., Mikaberidze, A., Mundt, C., Zhong, Z. & McDonald, B.A. (2016b). An improved method for measuring quantitative resistance to the wheat pathogen *Zymoseptoria tritici* using high throughput automated image analysis. Phytopathology, 106, 782–788.

Stowe, K.A. (1998). Experimental Evolution of Resistance in Brassica rapa: Correlated Response of Tolerance in Lines Selected for Glucosinolate Content. Evolution, 52, 703–712.

Teng, P.S. & Gaunt, R.E. (1980). Modelling systems of disease and yield loss in cereals. Agricultural Systems, 6, 131–154.

Thomas, M.R., Cook, R.J., & King, J.E. (1989). Factors affecting development of Septoria tritici in winter wheat and its effect on yield. Plant Pathology, 38, 246–257.

Waggoner, P. & Berger, R. (1987). Defoliation, disease, and growth. Phytopathology, 77, 393–398.

*Yates, S., *Mikaberidze, A., Krattinger, S.G., Abrouk, M., Hund, A., Yu, K., Studer, B., Fouche, S., Meile, L., Pereira, D., Karisto, P., McDonald, B.A. (2019). Precision phenotyping reveals novel loci for quantitative resistance to septoria tritici blotch,. Plant Phenomics, 3285904. (*equal contributions)

Zhan, J., Pettway, R. & McDonald, B. (2003). The global genetic structure of the wheat pathogen *Mycosphaerella graminicola* is characterized by high nuclear diversity, low mitochondrial diversity, regular recombination, and gene flow. Fungal Genetics and Biology, 38, 286–297.

Zhan, J., Stefanato, F. & McDonald, B. (2006). Selection for increased cyproconazole tolerance in *Mycosphaerella graminicola* through local adaptation and in response to host. Mol. Plant Pathol., 7, 259–268.

Zeller, M & Koella, J. (2017). The Role of the Environment in the Evolution of Tolerance and Resistance to a Pathogen. American Naturalist, 190, 389–397.

Zuckerman, E., Eshel, A. & Eyal, Z. (1997). Physiological Aspects Related to Tolerance of Spring Wheat Cultivars to Septoria tritici Blotch. Phytopathology, 87, 60–65.

